# A posteriori evaluation of molecular divergence dates using empirical estimates of time-heterogeneous fossilization rates

**DOI:** 10.1101/128314

**Authors:** Simon Gunkel, Jes Rust, Torsten Wappler, Christoph Mayer, Oliver Niehuis, Bernhard Misof

## Abstract

The application of molecular clock concepts in phylogenetics permits estimating the divergence times of clades with an incomplete fossil record. However, the reliability of this approach is disputed, because the resulting estimates are often inconsistent with different sets of fossils and other parameters (clock models and prior settings) in the analyses. Here, we present the λ statistic, a likelihood approach for *a posteriori* evaluating the reliability of estimated divergence times. The λ statistic is based on empirically derived fossilization rates and evaluates the fit of estimated divergence times to the fossil record. We tested the performance of this measure with simulated data sets. Furthermore, we applied it to the estimated divergence times of (*i*) Clavigeritae beetles of the family Staphylinidae and (*ii*) all extant insect orders. The reanalyzed beetle data supports the originally published results, but shows that several fossil calibrations used do not increase the reliability of the divergence time estimates. Analyses of estimated inter-ordinal insect divergences indicate that uniform priors with soft bounds marginally outperform log-normal priors on node ages. Furthermore, *a posteriori* evaluation of the original published analysis indicates that several inter-ordinal divergence estimates might be too young. The λ statistic allows the comparative evaluation of any clade divergence estimate derived from different calibration approaches. Consequently, the application of different algorithms, software tools, and calibration schemes can be empirically assessed.

## Introduction

Molecular clock analyses can be used to infer the geological origin of the most recent common ancestor (MRCA) of any two extant species or clades, even with an incomplete fossil record. This is of course very appealing to a broad range of scientists, including molecular biologists, systematists, and palaeontologists, who have extensively applied the molecular clock concept in their studies. The concept of the molecular clock was initially established empirically (Zuckerkandl and Pauling, 1962). Its theoretical foundations are based on the neutral theory of molecular evolution (Kimura, 1968; King and Jukes, 1969). Thus, the similarity of two given orthologous DNA sequences is expected to decay with an approximately constant rate over time. DNA sequence divergence can therefore be used to infer relative species divergence times. An absolute time scale is achieved by calibrating the relative age of a node with the absolute age of a fossil associated with the node. However, this simple approach has been subject to substantial criticism, because (*i*) calibration errors can accumulate (e.g.(Graur and Martin, 2004)) and (*ii*) substitution rates are not constant over time (Drummond, 2006). We now know that these are biologically unrealistic assumptions. This problem has been partially solved by the development and subsequent application of so-called relaxed clock methods, which do not assume a constant substitution rate among taxa and use multiple fossil calibrations. Relaxed molecular clock models not only offer an improved fit to the empirical data, but also are capable of accommodating seemingly contradictory calibration points, as well as a range of either explicitly or implicitly assumed prior probability distributions of node ages. Consequently, the application of relaxed molecular clocks became a standard procedure. The most commonly used software tools implementing relaxed molecular clock methods are BEAST (Drummond *et al*., 2007), MCMCtree (Yang, 2007), and dpp-div (Heath *et al*., 2012). All three software packages rely on a Bayesian approach to estimate and calibrate divergence times among clades. The reliability of the inferred estimates using a Bayesian approach strongly depends among other factors on a correct phylogenetic assignment of fossils (Parham *et al*., 2012) and proper choice of node age priors (Inoue *et al*., 2010; Warnock *et al*., 2012). Explicit *a priori* assignment of fossils and the choice of node age priors is not straightforward and is typically done in a highly subjective manner, resulting in substantial debate over the interpretation and utility of the results. At the same time, differences in the specification of analysis parameters typically also results in different divergence time estimates. The search for methods that can inform parameter choice is consequently of fundamental importance for further improving molecular tree calibrations. While currently accepted practice, arbitrary choice of node age priors can drastically affect results (Inoue *et al*., 2010; Warnock *et al*., 2012). The recently introduced fdpp-div (Heath *et al*., 2014) as well as other implementations of the fossilized birth-death process (Zhang *et al*., 2016) seek to substitute the arbitrary choice of fossilization rate distributions with the application of a model that estimates rates of speciation, extinction, and fossilization as well as the proportion of sampled extant species directly from the data. The approach assumes a constant fossilization rate within a given taxon across time. This, however, is unrealistic, because the fossil record documents that fossilization rates are not constant over time and across taxa. Thus, fdpp-div (Heath *et al*., 2014) and other approaches (Zhang *et al*., 2016) still rely on a palaeontologically unrealistic assumption. In this study, we introduce a new *a posteriori* approach which scores dated phylogenies for their congruence with the fossil record using an empirically derived time-dependent fossilization rate. The approach allows a critical *a posteriori* appraisal of dated phylogenies and in consequence the effect of node age priors on estimated divergence dates.

## New Approaches

It has been recognized in paleontology that the stratigraphic range of a taxon can be extended by taking into account evidence from pylogenetic studies. This resulted in the concept of a *ghost lineage*, which is defined as the geological time interval during which a clade is not documented by a fossil, but its is postulated based on the existence of a fossil from the clades sister taxon (Norell and Novacek, 1992) (Figure 1). Analogous to the notion of a *ghost lineage*, we introduce the notion of an *invisible lineage* as occupying the time interval over which the stratigraphic range of a clade is extended by molecular clock dating. Invisible lineages differ from ghost lineages in requiring data beyond the ages of fossils and their phylogenetic position. However, both ghost lineages and invisible lineages represent time intervals during which a taxon existed, but left no known fossil documentation. This absence of documentation is caused in both cases by one of three possibilities: (*i*) no fossil has been preserved, (*ii*) a fossil has been preserved, but relevant apomorphic traits have not been preserved, (*iii*) a fossil has been preserved, but relevant apomorphic traits had not yet evolved. The factors that influence (*iiii*) can be broadly categorized into taxon-specific (intrinsic) and taxon-unspecific (extrinsic) factors. Taxon-specific factors include all properties of organisms that affect the probability of any given individual of a taxon to be preserved as a fossil. This includes anatomical properties (e.g., an insect of larger size is less likely to be trapped in amber), but also ecological properties (e.g., an insect placing eggs in tree bark is more likely to be trapped in amber than an insect not doing so) and even sociological aspects (e.g., the number of paleoentomologists working on dipterans determines to a large extent the probability that a dipteran fossil will be described and phylogenetically placed). Taxon-unspecific factors apply to a taxonomically wide range of taxa and include the availability of sediments from a particular time range, the presence of LagerstϤtten, and diagenetic processes (see (Holland, 2016)). Taxon-unspecific properties also comprise sociological aspects, such as large regional differences in the investigation of fossil LagerstϤtten. The fossilization rates for a given time period can be inferred from the inverse of the average duration of all ghost lineages covering a particular time period within a phylogenetic tree. Invisible lineages are explained by the same fundamental process. Therefore, these empirically derived rates of fossilization can be used to evaluate estimates of clade origin by application of a (relaxed) molecular clock, and we can assume that the fossilization rates of ghost lineages, which we can infer from the data, are identical to those of invisible lineages. Such an approach incorporates information on taxon-unspecific properties of ghost lineages, but it is agnostic of the taxon-specific properties of these ghost lineages. The rationale is that taxon-unspecific properties mostly determine fossilization probabilities across taxa and, in consequence, the probability of identifying fossils of a particular clade in relation to its geological time of origin. It is therefore a time-dependent model. This assumption is of course only fully justified if phylogenetic analyses are taxonomically restricted, for example, to insects, birds, or mammals. Theoretically, this assumption can be relaxed if sufficient fossil data is available. However, there is currently not enough data to construct a robust time-heterogeneous model, which could potentially deal with taxon-specific <u>and</u> -unspecific properties.

**FIG. 1.**
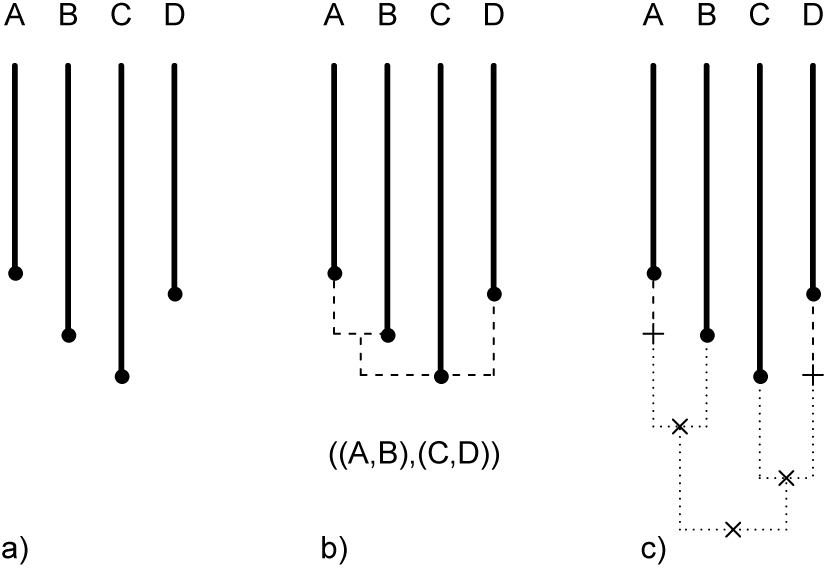
Graphical illustration of the definition of the terms *ghost lineage* and *invisible lineage*. a) Earliest fossil documentation of taxa A, B, C, and D. b) *Ghost lineages* (dashed lines) inferred from considering the fossil data in conjunction with the phylogenetic relationship between the taxa. c) *Invisible lineages* (dotted lines) inferred from considering the fossil data, the phylogenetic relationship between the taxa, and a molecular dating hypothesis on the age of nodes.

Our time-dependent model is constructed in the following way. If *G* is the set of ghost lineages, then for each *g*∈*G* we define *a_g_* ≤ *b_g_* as the starting times and end times of each ghost lineage. In order to derive a conservative estimate, *b_g_* represents the oldest possible appearance of the older fossil and *a_g_* the youngest possible appearance of the younger fossil defining the ghost lineage. We then derive a function λ(*t*) for all non-empty *G_t_*:

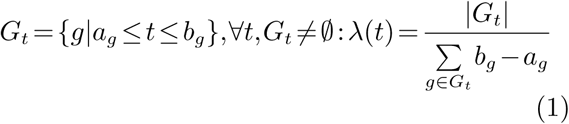

Thus, λ(*t*) is the inverse of the mean length of all ghost lineages spanning any point in time *t*. The number of ghost lineages spanning *t* determines the reliability of the statistic. Λ(*t*) can be interpreted as the fossilization rate at *t*. Higher values of λ(*t*) indicate shorter ghost lineages and correspond to conditions more favorable to fossilization. This implies that the fossil record has a higher time resolution and the fossil record at time *t* is becoming more informative. For a point in time not covered by at least one interval, the function is undefined. More formally, λ(*t*) is best understood as part of a time-heterogeneous exponential distribution, giving a probability *p* that the length of ghost lineages starting at time *t* exceed a length of *x* as

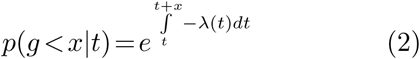

If the conditions that affect fossilization are stationary over time, then the distribution of all ghost lineages would be exponential with a parameter λ = λ(*t*).

The model introduced here provides a view on the quality of the fossil record using the fossil record itself. Next, we compare it to the implicit model presented by a molecular dating hypothesis. Given a fossil representative at time *T_f_* and a molecular estimate for the same node at time *T_m_*, we will have an observed invisible lineage Δ*T* = *T_m_* – *T_f_*. The likelihood *f* of observing a difference this large or larger is given by

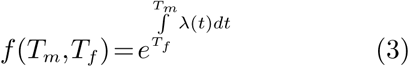
which is a straightforward result from (1). We define a fit score *S* of each node as

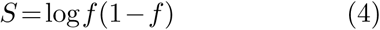
which equally treats both the probability of an invisible lineage larger than Δ*T* and one smaller than Δ*T*. Finally, the total fit score for a dating hypothesis is the sum over all nodes for which *S* has been calculated. We call this the λ-score of a tree calibration. A higher λ-score indicates a higher degree of consistency of the data with the fossil record. This method improves on other a posteriori approaches (e.g., (Near *et al*., 2005)) in two ways. Firstly, it weights the gaps between the molecular date and the oldest fossil representative depending on the time resolution of the fossil record and secondly, it penalizes gaps that are smaller than the fossilization rate suggests.

## Results

Analyzing simulated data, we first confirmed that the λ statistic is able to improve dating. In order to generate data sets with properties representative of empirical data, we used different mean fossilization rates covering the assumed range of empirical fossilization rates, different extent of evolutionary rate heterogeneity in the molecular data, and different numbers of taxa. Using these data sets, we calibrated trees considering all possible combinations of simulated fossil calibration points (see details in Material and Methods). Depending on the mean fossilization rate, the rate heterogeneity of the molecular sequence data, the different numbers of taxa and the different fossil calibration points and node age priors, the estimated node ages varied, in some instances extensively. Among the calibrations, we identified the one with the smallest sum of absolute deviations of the simulated (true) and estimated node ages. The results show that the λ statistic is a powerful measure to help at identifying the calibration scheme with the smallest deviation (the smallest sum of absolute differences). In cases in which the calibration scheme with the smallest deviation is not the best scheme chosen by the λ-score, errors due to selecting a suboptimal calibration are small (Table 1, Figure 2). In several instances, the calibration with the closest match to the true (simulated) node ages contained at least one node estimated to be younger than the corresponding fossil. These calibrations are partially inconsistent with the fossil record and a λ-score can not be calculated for them. If the inconsistent calibrations are removed, the percentages of correctly identified optimal calibrations increased, while the time differences between estimated and true node ages decreased (Table 1, Figures 2 and 3).

**FIG. 2.**
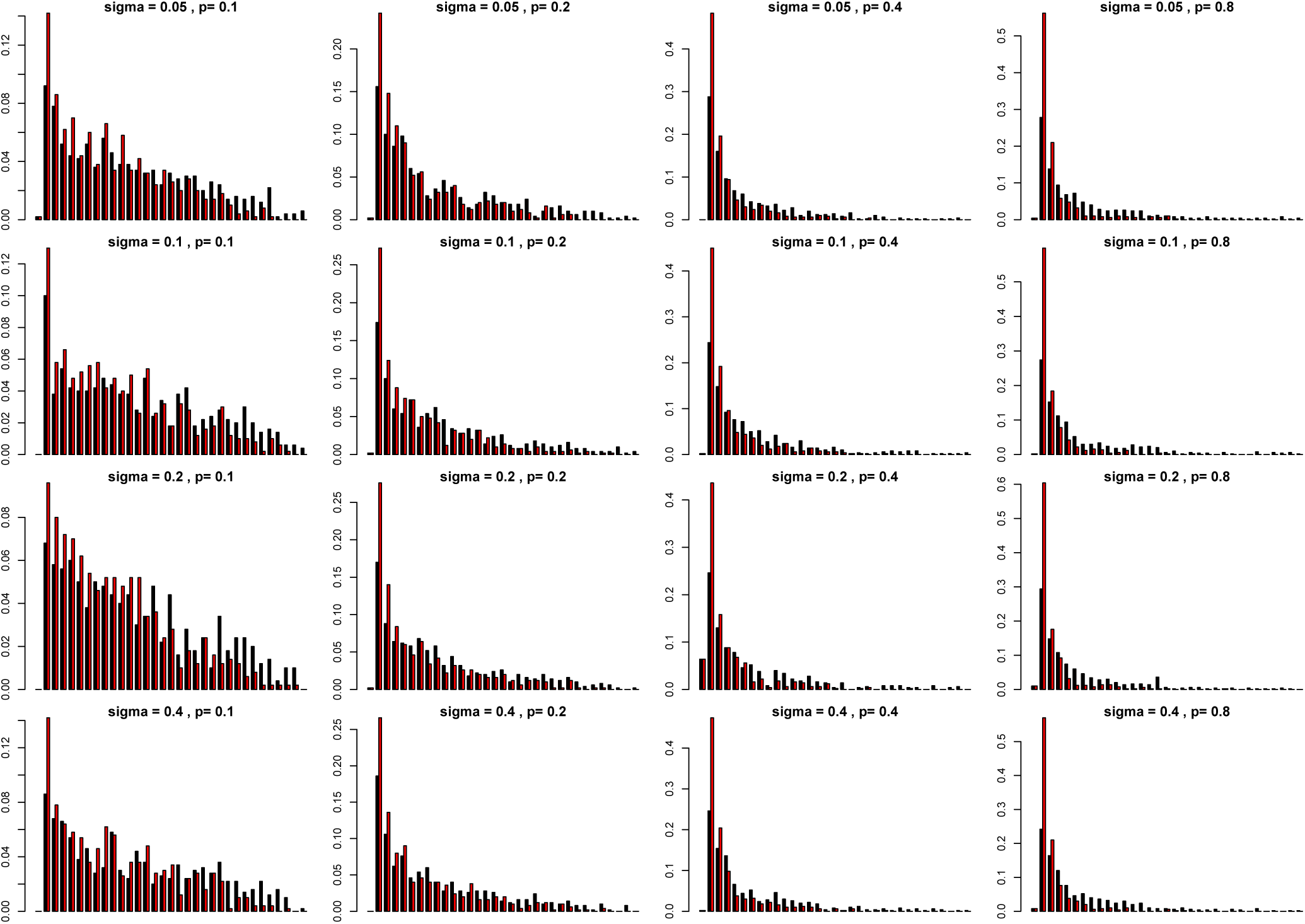
Frequency distribution of ranks of the dating hypotheses with the highest λ-scores for seven taxa with simulated λ(*t*). Black bars show ranks when all hypotheses are considered, red bars show ranks when only trees with no node younger than the oldest fossil are considered.

**FIG. 3.**
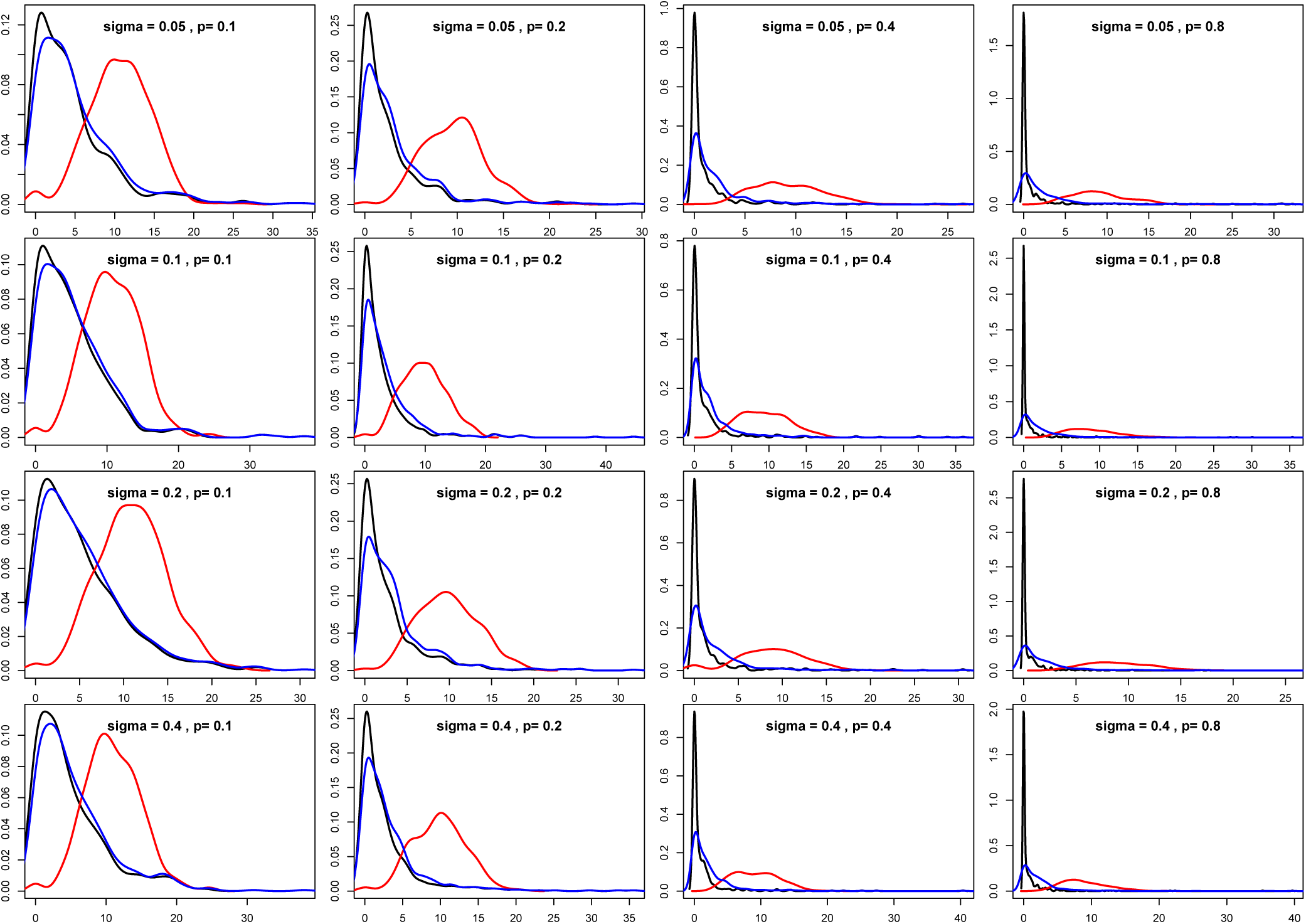
Density curves of the time differences between trees and the tree matching the true divergence dates the closest for seven taxa with simulated λ(*t*). Red line: random tree. Blue line: tree with the highest λ-score, taking all trees into consideration. Black line: tree with the highest λ-scores ignoring trees with nodes dated to younger ages than the oldest fossil.

**Table 1.** Recovery rate of the best tree calibration using the λ-score in simulations. *N* gives the number of taxa in the analyzed phylogeny, *p* and *σ* the quality of the fossil record and the parameter determining the departure of branch length standard deviations away from a clock like tree, respetively. # gives the number of replicates for the simulation setup in 100s. If empirical data from the fossil record of insects were used, *p* is not applicable (NA). *R* gives the proportion of best dating hypotheses found λ(*t*) statistics. *t_lost_* indicates the mean time per node, by which the tree with the highest λ-score is worse than the tree closest to the real divergence dates. 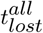 gives this time for a dating hypothesis chosen at random rather than with the aid of the λ-score and thus acts as a control. *R*′ and 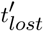 give the values if the trees under consideration are removed from any trees in which at least one node is dated as younger than the oldest fossil. *T_R_* gives the age assigned to the root of the tree in the simulations in 100 Ma.

| $N$ | $p$ | $\sigma$ | $\#$ | $R$ | $R'$ | $t_{lost}$ | $t'_{lost}$ | $t_{lost}^{all}$ | $T_R$ |
| --- | --- | --- | --- | --- | --- | --- | --- | --- | --- |
| 7 | 0.8 | 0.05 | 5 | 0.34 | 0.66 | 1.55 | 0.24 | 8.97 | 1 |
| 7 | 0.4 | 0.05 | 5 | 0.34 | 0.57 | 1.29 | 0.50 | 9.12 | 1 |
| 7 | 0.2 | 0.05 | 5 | 0.23 | 0.34 | 1.90 | 1.34 | 9.54 | 1 |
| 7 | 0.1 | 0.05 | 5 | 0.16 | 0.23 | 3.53 | 2.96 | 10.61 | 1 |
| 7 | 0.8 | 0.10 | 5 | 0.33 | 0.67 | 1.65 | 0.21 | 8.78 | 1 |
| 7 | 0.4 | 0.10 | 5 | 0.30 | 0.55 | 1.39 | 0.50 | 9.36 | 1 |
| 7 | 0.2 | 0.10 | 5 | 0.23 | 0.36 | 2.11 | 1.40 | 9.77 | 1 |
| 7 | 0.1 | 0.10 | 5 | 0.16 | 0.20 | 3.69 | 3.25 | 10.70 | 1 |
| 7 | 0.8 | 0.20 | 5 | 0.34 | 0.67 | 1.59 | 0.25 | 9.03 | 1 |
| 7 | 0.4 | 0.20 | 5 | 0.34 | 0.54 | 1.36 | 0.48 | 8.72 | 1 |
| 7 | 0.2 | 0.20 | 5 | 0.23 | 0.38 | 1.99 | 1.31 | 9.82 | 1 |
| 7 | 0.1 | 0.20 | 5 | 0.11 | 0.16 | 3.93 | 3.53 | 10.93 | 1 |
| 7 | 0.8 | 0.40 | 5 | 0.30 | 0.68 | 1.74 | 0.22 | 8.91 | 1 |
| 7 | 0.4 | 0.40 | 5 | 0.31 | 0.56 | 1.47 | 0.44 | 9.21 | 1 |
| 7 | 0.2 | 0.40 | 5 | 0.24 | 0.34 | 1.76 | 1.24 | 9.84 | 1 |
| 7 | 0.1 | 0.40 | 5 | 0.15 | 0.21 | 3.71 | 3.18 | 10.94 | 1 |
| 7 | NA | 0.20 | 10 | 0.35 | 0.54 | 14.17 | 7.97 | 44.70 | 3 |
| 7 | NA | 0.40 | 10 | 0.37 | 0.54 | 13.29 | 7.78 | 45.82 | 3 |
| 9 | 0.4 | 0.10 | 5 | 0.16 | 0.33 | 1.61 | 0.86 | 8.89 | 1 |
| 11 | 0.4 | 0.10 | 5 | 0.05 | 0.19 | 2.51 | 1.20 | 9.22 | 1 |

The degree to which branch lengths of trees were made non-clock-like (simulation parameter *σ*) did not alter the reliability of the λ statistic for identifying the best (calibrated) tree (Table 1). However, the choice of fossilization rates does affect results. With higher fossilization rates, implying that the fossil record is more informative, the highest λ-scoring trees were more often the *a priori* known optimal trees (Table 1). Even if the fossilization rates were low, implying that the fossil record is less informative (see Material and Methods), 11% of the best λ-scoring calibrations were the optimal ones, compared to 3% when randomly picking from the calibrations. With the most favorable choice of parameters, the rate of recovery of the optimal calibration increased to 34%. This implies that an increase in the mean quality of the fossil record increased the rate at which optimal calibrations were identified, and decreased the time difference between estimated and true node ages. The best λ-scoring calibrations had an average error per node less than 4 Ma greater than the calibrations with the least deviations in the simulations with the lowest fossilization rate and less than 2 Ma when analyzing data sets simulated with the highest fossilization rate (for details see Material and Methods,Table 1, Figure 3), while the mean error of all calibrations regardless of λ-score was about 10 Ma in all cases. The error from suboptimal calibration choice therefore decreased by 6080% when the λ-score was used. Optimal calibrations are recovered in 30 % of cases when analyzing trees with seven terminal taxa, 16 % of trees with nine terminals and 4.8% of trees with eleven terminals. The absolute time difference per node increases from 1.39 Ma to 1.6 Ma and 2.5 Ma, respectively, while calibrations not chosen through the λ-score showed on average 9.4 Ma, 8.9 Ma and 9.2 Ma absolute time differences between estimated and true node ages. After removing calibrations inconsistent with the fossil record, the recovery rate of the optimal calibrations increased to 55% when analyzing seven, 33% when analyzing nine and 19% when analyzing eleven taxa, while the absolute time differences between estimated and true ages per node dropped to 1.4Ma, 0.9 Ma and 1.2 Ma, respectively. It should be noted that the chances of a randomly chosen calibration being the optimal one are reduced from 3% (seven taxa) to 0.7% (nine taxa) and 0.2% (eleven taxa), respectively. The simulations show that a larger number of terminal taxa decreases the chance of recovering the optimal calibration, but the chance to identify the optimal calibration decreases to a lesser degree than the increase in total number of trees from which the optimal one has to be picked (Figure 4).

**FIG. 4.**
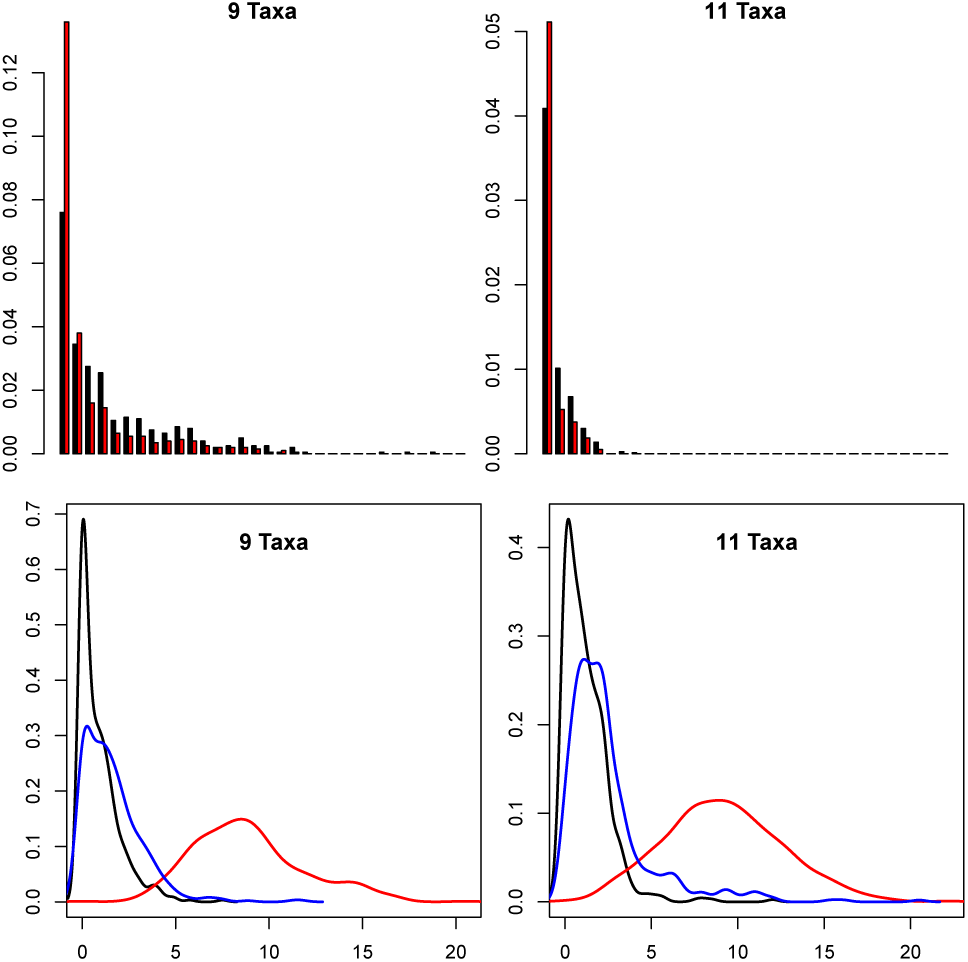
Frequency distributions and density curves for larger numbers of taxa. Top row: frequency distributions of the ranks of the dating hypotheses with the highest λ-scores for nine and eleven taxa with simulated λ(*t*). Black bars show ranks when all hypotheses are considered, red bars show ranks when only trees with no node younger than the oldest fossil are considered. Bottom row: density curves of the time differences between trees and the tree matching the true divergence dates the closest when analyzing nine and eleven taxa with a simulated λ(*t*). Red line: random tree. Blue line: tree with the highest λ-score, taking all trees into consideration. Black line: tree with the highest λ-scores, ignoring trees with nodes dated to younger ages than the oldest fossil.

Before we tested the λ statistic with empirical data, we first inferred an empirical λ(*t*) for studying insects using 829 pairs of adelpho (sister) taxa from PaleoDB (Kiselev *et al*., 2014). The resultant λ(*t*) curve shows major changes of fossilization rates roughly correlated with era changes in the Phanerozoic. During the Paleozoic, λ(*t*) peaks of fossilization rates correlate with well known fossil Lagerstätten. The largest value is reached at 253 Ma and results from the high frequency of putative sister taxa described from a single locality (Warners Bay, New South Wales, Australia (Tillyard, 1926)). In the Mesozoic, values of λ(*t*) decrease. While some localities give rise to local peaks in λ(*t*), they are far less pronounced compared with the Paleozoic. In the Cenozoic, we find another increase of λ(*t*) with peaks associated with conservation deposits, such as the Baltic amber. The resultant values of λ range from 0.017 Ma^−1^ to 0.2 Ma^−1^, with a mean of 0.045 Ma^−1^. These values are smaller than in all simulated data sets (see Figure 5 for used time intervals and the resultant time series of λ(*t*)). In order to test the influence of the empirically derived λ(*t*) on the ability of the λ statistic to detect optimal trees, we substituted the simulated function λ(*t*) with the empirically derived insect-specific λ(*t*) scores in our simulated data sets and repeated the series of tree calibrations. Using a 7-taxon tree and the empirical insect-specific λ(*t*) scores, the optimal tree calibrations are recovered in 37% of the replicates. Next, we increased the age of the root of the simulated trees to use the full potential of the empirical insect-specific λ(*t*) scores. The best λ-scoring calibrations had a mean age difference of 13.3 Ma per estimated node to the true node ages compared with 45.8 Ma difference for a randomly selected calibration (Figure 6). After removing inconsistent calibrations, optimal calibrations were identified in 53.8% of the replicates and the mean absolute time difference between estimated and true node ages dropped to 7.7 Ma (Table 1, Figure 6). In conclusion, these results show that the λ statistic can be used to select optimal calibration schemes based on empirical data sets. Additionally, the analysis should be limited to nodes at which all compared trees are consistent with the fossil record.

**FIG. 5.**
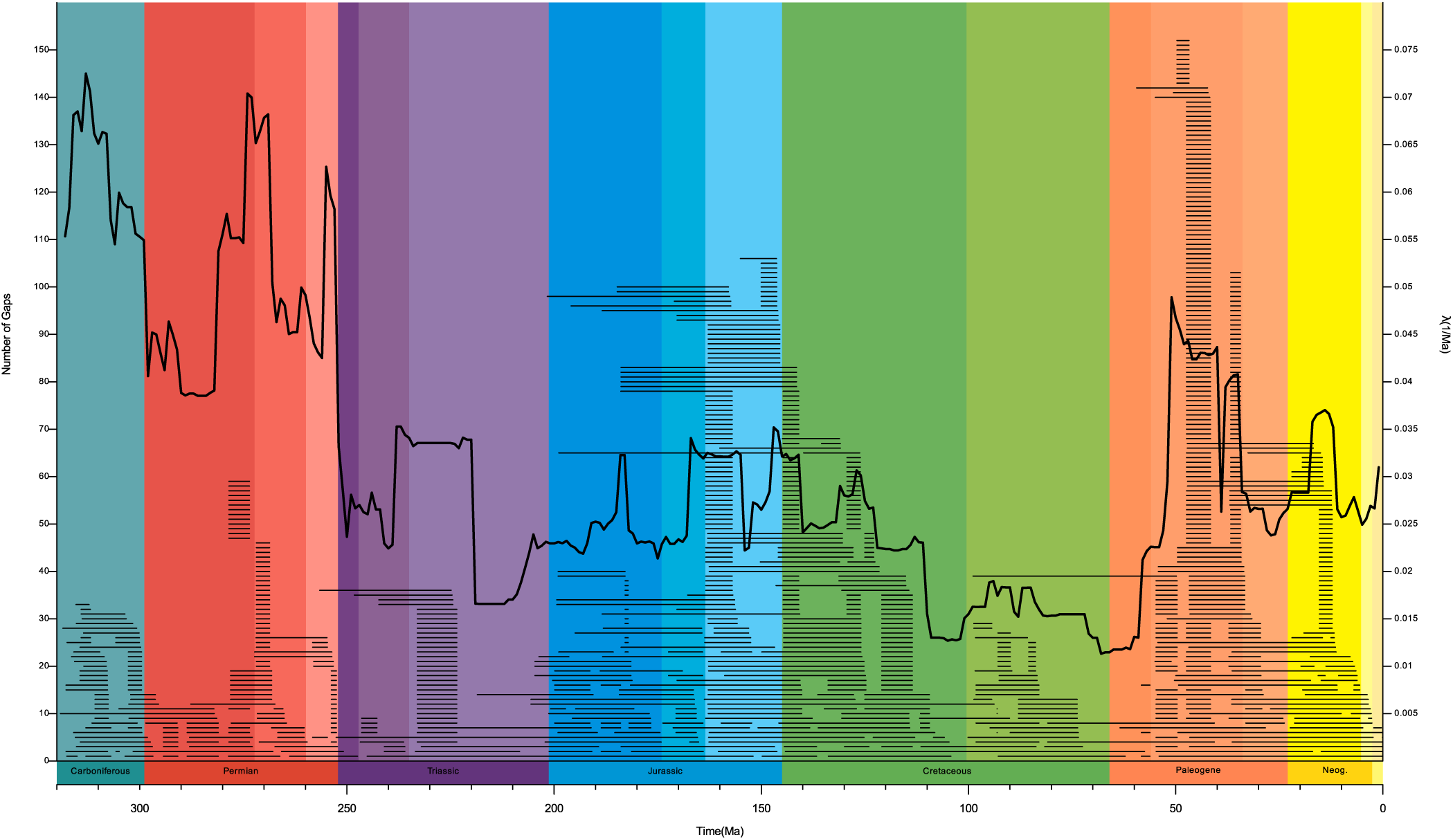
The insect-specific λ(*t*) through deep time. The gaps between first appearance dates used to calculate the curve are indicated by horizontal lines.

**FIG. 6.**
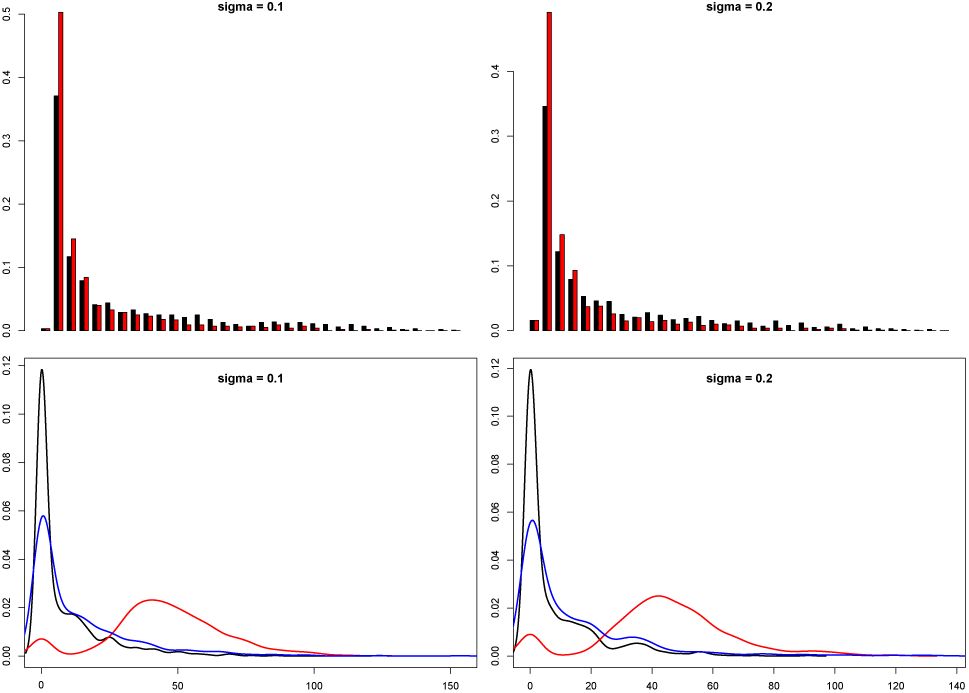
Frequency distributions and density curves for the empirical λ(*t*). Top row: frequency distributions of the ranks of the dating hypotheses with the highest λ-scores when analyzing seven taxa using an insect-specific λ(*t*). Black bars show ranks when all hypotheses are considered, red bars show ranks when only trees with no node younger than the oldest fossil are considered. Bottom row: density curves of the time differences between trees and the tree matching the true divergence dates the closest when analyzing seven taxa using an insect-specific λ(*t*). Red line: random tree. Blue line: tree with the highest λ-score, taking all trees into consideration. Black line: tree with the highest λ-scores, ignoring trees with nodes dated to younger ages than the oldest fossil.

Following the evaluation of the λ statistic with simulated data, we used the λ statistic to evaluate two published tree calibrations based on empirical molecular sequences and fossil data. We first evaluated a published tree calibration of the beetle supertribe Clavigeritae (Coleoptera: Staphylinidae: Pselaphinae) (Parker and Grimaldi, 2014). This data set was chosen, because one fossil used in the calibration was used in two variant calibrations by the authors who initially published the data (Parker and Grimaldi, 2014). In such a situation, the λ statistic should help to identify the tree calibration that best fits the fossil record. We used a selection of all possible subsets of fossil calibration points provided by Parker and Grimaldi (Parker and Grimaldi, 2014) to check whether or not the inclusion or exclusion of fossil calibration data changed the results of the tree calibration (details in Material and Methods). With this approach, we generated 255 calibrated trees with a wide range of λ-scores. Some subsets of tree calibrations yielded age estimates younger than the oldest fossil representative and these trees were subsequently not considered. Some calibration node subsets yielded clusters of similar λ-scores, which correlate with very similar node age estimates (Figure 7). These particular subsets share four fossil calibration points and are different in all possible combinations of presence and absence of the other four fossil calibration points. The tree calibration with the highest λ-score is very close to the original published result (Figure 8), supporting the idea of an origin of Clavigeritae in the Upper Cretaceous. Most importantly, the λ statistic showed that the inclusion of *Protoclaviger trichodens* (Parker and Grimaldi, 2014), the only representative of the in-group and controversial placement of the non-described “Fossil A” (Arhytodini), have no notable impact on the tree calibration even when used in a non-conservative manner (as in analysis #5 conducted by (Parker and Grimaldi, 2014)). In a second analysis, we compared contradicting tree calibrations of a large insect data set (Misof *et al*., 2014; Tong *et al*., 2015). In this analysis, we (*i*) compared the distribution of node age estimates among tree calibrations based on separate data partitions and (*ii*) evaluated the fit of the different published tree calibration approaches to the fossil record (see Material and Methods for details). Applying the λ statistic, we found that the calibration scheme applied by Tong *et al*. (Tong *et al*., 2015) that rests on uniform priors with soft bounds scored higher than the original calibration applied by Misof *et al*. (Misof *et al*., 2014), who used log-normal priors. However, Tong *et al*. (Tong *et al*., 2015) favored a tree calibration using soft minima. This scheme showed inferior λ-scores compared with those obtained when using log-normal priors as done by Misof *et al*. (Misof *et al*., 2014) (Figure 9). The inclusion of an additional calibration point from a roachoid fossil reduced the λ-score even more, but it is unclear what its impact would have been if it was included in an analysis using log-normal or pseudo log-normal priors for other nodes. The consensus dating hypothesis based on the ten best scoring meta-partitions of all four different calibration approaches (Figure 10) corroborated most of the results presented by Misof *et al*. (Misof *et al*., 2014). With our approach, the origin of Hexapoda is dated to 480 Ma, that of Insecta to 445 Ma, that of Pterygota to 400 Ma, and that of the origin of Holometabola to 340 Ma. The clade containing Orthoptera and Blattodea, crucial to the discussion of the roachoid fossil, is dated to an age slightly younger than the onset of the Triassic. The origin of extant Polyneoptera is estimated at 290 Ma, which is about 10 Ma younger than suggested by Misof *et al*. (Misof *et al*., 2014). The radiation of parasitic lice is dated to 59 Ma, remaining post-Cretaceous as suggested by Misof *et al*. (Misof *et al*., 2014).

**FIG. 7.**
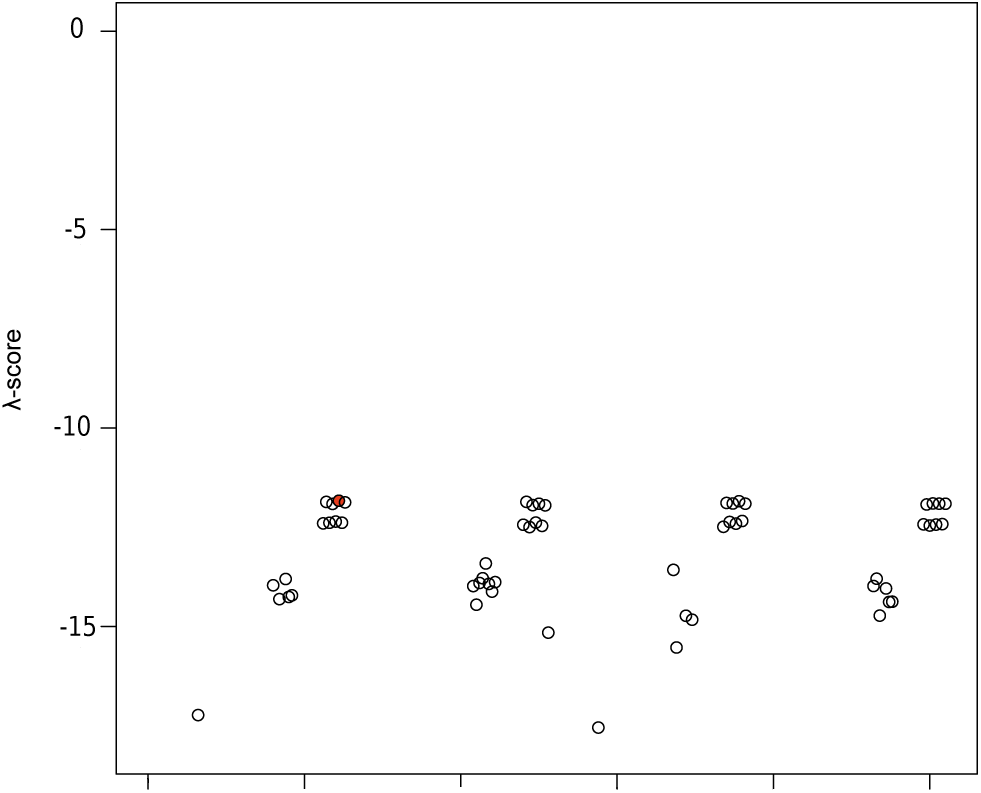
λ-scores obtained from 255 alternative calibration schemes used to study the divergence times of Clavigeritae. The filled circle indicates the maximal value.

**FIG. 8.**
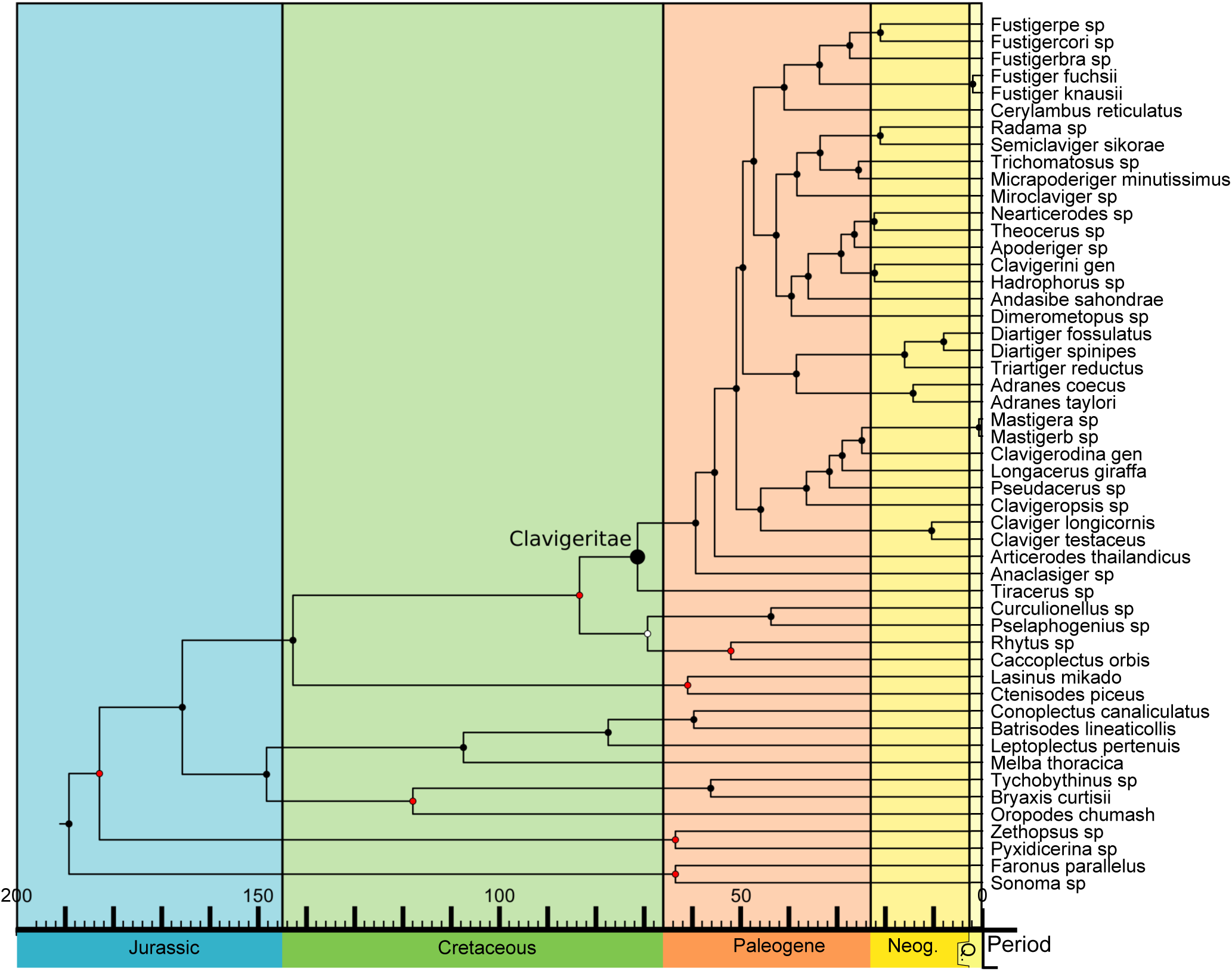
Dated phylogeny for Clavigeritae, with dates taken from the calibration with the highest λ-score.

**FIG. 9.**
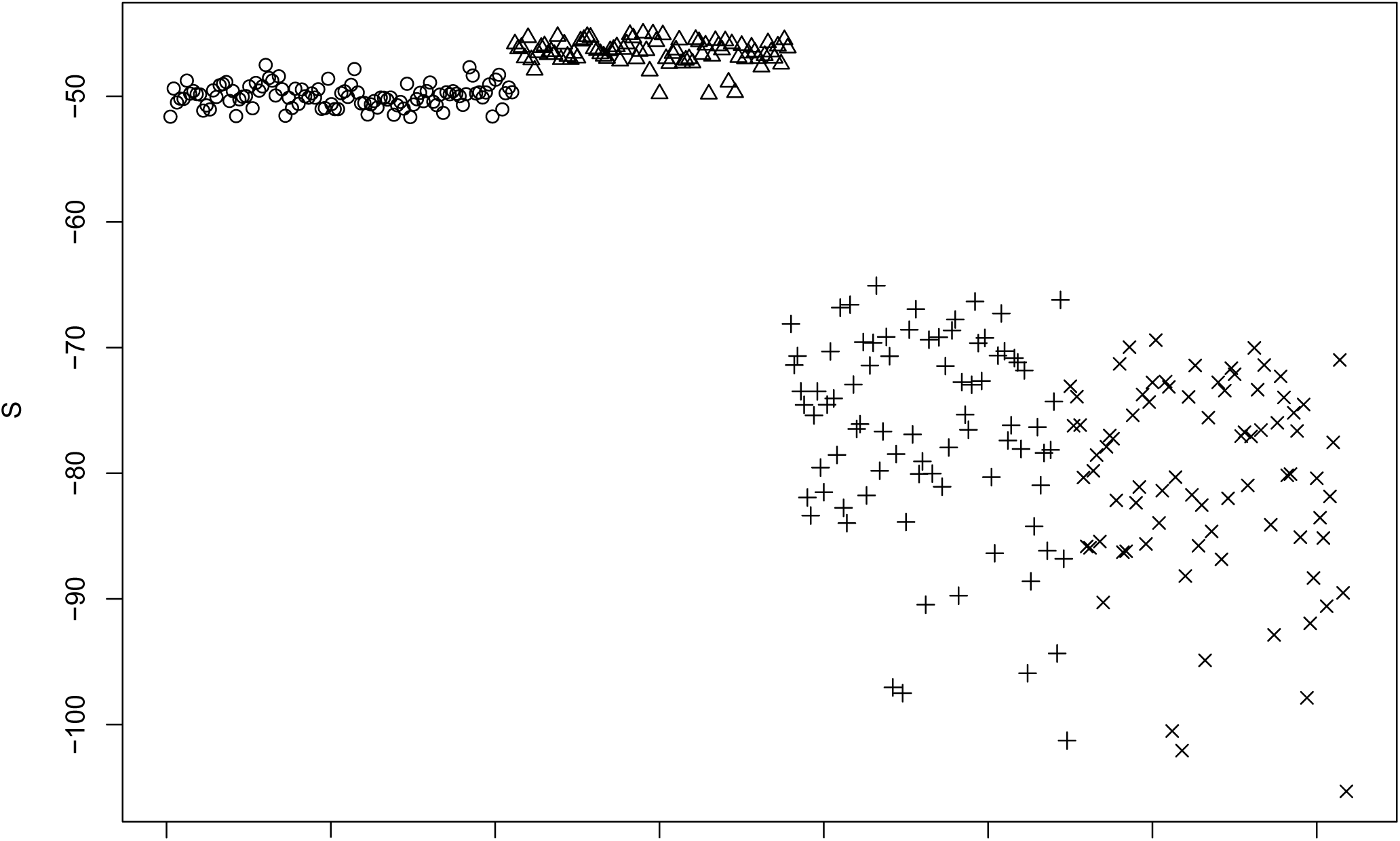
λ-scores calculated for the meta-partitions of the insect dataset. Circles: Calibration scheme used by Misof *et al*. (Misof *et al*., 2014); triangles: first calibration scheme used by Tong *et al*. (Tong *et al*., 2015), pseudo-log-normal; crosses: second calibration scheme used by Tong *et al*. (Tong *et al*., 2015), soft minima only; xs: third calibration scheme used by Tong *et al*. (Tong *et al*., 2015), soft minima with additional calibration point.

**FIG. 10.**
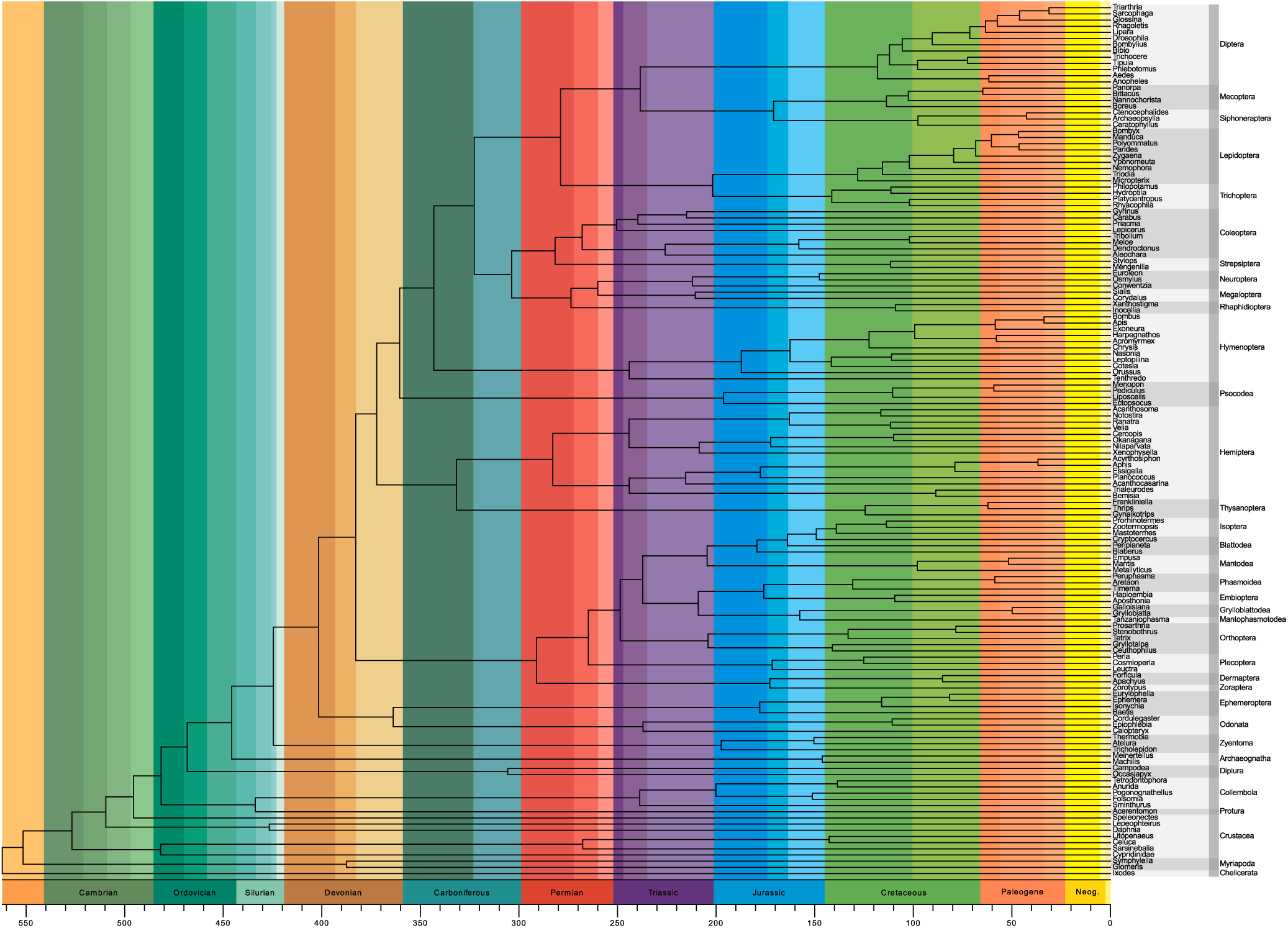
Consensus dating hypothesis obtained for the insect tree when considering the ten meta-partitions with the highest λ-scores. All of the highest scoring meta-partitions were calibrated using the pseudo log-normal scheme suggested by Tong *et al*. (Tong *et al*., 2015).

## Discussion

Divergence time estimates based on tree calibrations with too small a number of calibration points, missing or unusually small error bars have been vigorously criticized as unscientific story telling (4). Besides the large error rate of tree calibrations, which are often ignored, the uncertain taxonomic placement of fossil calibration points and the arbitrary choice of node age priors are the major targets of criticism. At its present state of methodology, results of tree calibrations can only be scientifically tested with the admittedly often uncertain taxonomic placement of relevant fossils. Using the here introduced λ-score, which measures the fit of a tree calibration to the fossil record, we aim to identify the optimal tree calibration. Relying on simulated data, we demonstrate the utility of this *a posteriori* approach. If the optimal tree calibration was missed, the absolute time difference in mean node age estimates between the optimal and the chosen tree calibration was small compared to the mean across all tree calibrations. Since the λ-score negatively correlates with errors of node age estimates, the fossilization rate function λ(*t*) can be used to make informed choices of node age priors in tree calibrations based on a Bayesian approach. This is a major advance in tree calibration studies. An empirical fossilization rate function λ(*t*), which is a cornerstone of our approach, can be derived with the data archived in databases such as PaleoDB (Kiselev *et al*., 2014). We compared such an empirically derived insect-specific λ(*t*) statistic with one that is based on simulated data to show that even in the case of higher mean values of the insect-specific λ(*t*) scores compared with the simulated ones, the λ-score is a reliable measure for identifying optimal or nearly optimal tree calibrations. However, the informative value of the λ-score decreases with a sparse fossil record. While databases like PaleoDB (Kiselev *et al*., 2014) or the insect-specific EDNA database (Mitchell, 2013) record taxonomic ranks of fossils, they do not provide data on the phylogenetic placement of the fossils. As a result, only a fraction of known fossils can be used to derive an empirical λ(*t*). Extensive phylogenetic work could solve this deficiency. Such analyses could also help to solve another problem that we observed when analyzing data from PaleoDB (Kiselev *et al*., 2014): the empirical insect λ(*t*)shows remarkably high values in the Paleozoic. This bias is likely a consequence of the comparatively small number of researchers who have studied paleozoic Lagerstätten and described new genera containing at least two species. Yet, despite these taxon-unspecific effects, we can show that the empirically derived fossilization rate function λ(*t*) works very well when applied to tree calibrations. The λ-score optimization rests on the assumption that fossilization rates are uniform across the analyzed taxa within a time slice. This assumption does not necessary hold and has the potential of introducing error, which should not be naively ignored. By comparison, fdpp-div citefdpp does resolve differences in fossilization rates among taxa, but can not resolve changes in fossilization rate over time. A possible extension of the λ statistic could therefore be to divide the taxon sample into taphonomic classes and calculate a class-specific λ(*t*). The major limiting factor for such class-specific λ(*t*)is the current availability of a sufficiently informative fossil record. Both approaches have their validity, depending on the taxonomic problem. In some taxa, changes in fossilization rate over time are likely negligible, while in others, changes in fossilization rate over time are expected to be large. Our reassessment of the Clavigeritae data set (Parker and Grimaldi, 2014) revealed that additional fossil calibration points, even if initially considered important, do not necessarily improve divergence time estimates. Futhermore, redundant soft minimum ages do not automatically decrease the accuracy of the divergence time estimates. This result suggests that minimum age constraints on node ages are relatively safe to use, even if the fossils are not well constrained in time, provided the time estimate is chosen at the youngest possible age. It is important to note that the only fossil calibration point placed within the Clavigeritae did not alter the tree calibration results. Thus, fossil calibration points of outgroups can in some instances be sufficient (and even better suited) to provide a solid framework for the application of relaxed clock models. This result also means that the choice of outgroup taxa can bias tree calibration significantly and therefore should be done with similar care as the choice of ingroup taxa. The reanalysis of insect order divergence times (Misof *et al*., 2014; Tong *et al*., 2015) demonstrates the power of the λ statistic to evaluate different calibration schemes. We show that the λ-score can be used to select optimal data partitions for calibration of trees. Applying the λ-score, we demonstrate that the calibration used by Tong *et al*. (2015) using soft uniform priors instead of log-normal priors produced higher λ-scores. The soft uniform priors assign more weight to slightly older ages, even if the 95 % confidence intervals are almost identical with those obtained by Misof *et al*. (2014). Therefore, in this case log-normal priors have a slight tendency to be too restrictive. Yet, a tree based on the best fitting meta-partitions from the entire range of analyses conducted by both Misof *et al*. (2014) and Tong *et al*. (2015) differed only in details and was very close to the original result presented by Misof *et al*. (2014). We observed a strong decrease in λ-scores when applying calibration schemes that use soft minima instead of log-normal priors favored by Tong *et al*. (2015). The tree calibration derived with soft minima and including the roachoid calibration shows the worst λ-score of all four available dating schemes. The node age estimates inferred by Misof *et al*. (2014) were often younger than the ones delivering a λ maximum for any particular node, but the estimated node ages inferred by Tong *et al*. (2015) using only soft minima were usually far older. Apparently, soft minimum priors are too permissive in this case. Our results do not allow us to fully address the suitability of the additional fossil considered by Tong *et al*. (2015) for calibrating the phylogeny of extant insect orders, since the use of soft minima already decreased the λ-score. However, we observed a slight decrease in λ-scores when including the roachoid fossil in our analyses of the dataset as compared to the ones not including it. Priors are effectively hypotheses on the relationship between clade age and the age of their oldest fossil representatives. Priors are thus hypotheses on the quality of the fossil record. The exclusive reliance on soft minima represents a very pessimistic view of the quality of the fossil record, which might be ill advised. With the λ statistic, we provide the means to empirically evaluate hypotheses on the quality of the fossil record of a particular taxon and thus pave the road to more objective tree calibrations in molecular systematics.

## Material and Methods

We used simulated data to demonstrate the validity of the λ(*t*)-score approach proposed herein. Furthermore, we evaluated the performance of this approach using two empirical data sets.

### Simulated data

Evolver (version 4.7a of the PAML package (Yang, 2007)) was used to generate three random rooted tree topologies with seven, nine, and eleven terminal taxa, respectively. A custom script in R (R Core Team, 2012) was used to generate node ages. The root node age was fixed to 100 Ma with the terminal nodes at 0 Ma. All other nodes ages were set by drawing from a uniform distribution constrained to be younger than the next ancestral nodes. λ(*t*) was simulated in the time range from 100 to 0 (today) Ma as a step-function, using uniform distributions on [0,*p*] to generate values for time intervals of 1 Ma duration. We ran simulations with *p* chosen from {0.1, 0.2, 0.4, 0.8}, thereby comparing variable fossilization rates across time periods. Increased values for *p* generate fossilization rates of higher maximum amplitude. Fossil calibration points were simulated in the following way: a set of fossil calibration points was created by generating a uniform variable *X* on [0, 1] and then using the divergence time *T* and λ(*t*) to generate a fossil for each node using (3) by setting *p*(*T,T_f_*) = *X* and solving for *T_f_*. This process can produce an age of 0 or less, creating an absence of fossils from the lineage. A molecular dataset of length 90,000 nucleotides for 7, 9 or 11 was created using Evolver under the Jukes-Cantor model. Branch lengths were defined as node age differences divided by 100 Ma, so that the sum of all branch lengths from the root to the tip were equal to 1. This tree was still clock like. In order to add stochastic deviations from a strict clock data set, we added a random number drawn from a normal distribution (*μ*= 0) to each branch length. The whole simulation procedure has two free simulation parameters, namely *p*, which allows one to adjust the quality of the fossil record and the standard deviation *σ*, used to specify the variability of the branch lengths and the deviation from a strict clock like tree. Altogether, we generated 500 trees and datasets for each of the 16 simulation set ups with *p* ∈ {0.1,0.2,0.4,0.8},*σ* ∈ {0.05,0.1,0.2,0.4}. Each simulated dataset was used to calibrate the corresponding known tree with the Bayesian approach implemented in MCMCtree (Yang, 2007) using each possible subset of fossil calibration points (implemented as soft minimum ages). Results of the tree calibrations were scored based on the generated λ(*t*) using the score *S* as defined in (4). To check for the effect of tree size, a single run of 500 replicates using *p* = 0.4 and *σ* = 0.1 was performed for topologies with nine and eleven terminal taxa. The number of different combinations of calibration points *K* depending on the number of taxa *N* in the tree is given by *K* = 2*^N^*^−2^ – 1. Simulation setups and their parameters are listed in Table 1. In two simulation setups, we did not use a simulated function λ(*t*), but a single λ(*t*) inferred from empirical data. The empirical dataset used to infer an insect-specific λ(*t*) was generated by using information stored in PaleobioDB (Kiselev *et al*., 2014). We first downloaded the complete data on the fossil record of insects from the database. We then selected all genera that contained precisely two fossil species, which were subsequently considered putative sister taxa. For both fossils, we recorded the earliest appearance in the fossil record (see Supplementary file for a list of taxa used). This dataset was then used to calculate λ(*t*) using (1). We simulated 1,000 replicates with the same topology that was used to analyzed the seven taxon trees with *σ* ∈ {0.2,0.4} and a root age of 300 Ma, assigning ages and fossil calibration points in the same way as in the first set of simulations. We increased the root age in these simulations to be able to utilize the full time range of the empirical insect λ(*t*). From each MCMC tree dating result (DR), we calculated the λ-score as well as the sum of absolute differences between estimated and real node ages. All DRs were subsequently ranked according to the absolute differences between estimated and real node ages. (i.e., a DR with a rank of 1 is the DR exhibiting the smallest deviations from the true ages). The best DR according to the λ-score is not always ranked best in terms of absolute differences between estimated and real node ages. Therefore, the difference in estimated node ages between the best λ-scoring calibration and the calibration with the smallest total deviation from the true divergence dates was also calculated. This measure allowed us to track by how much the best λ-scoring calibration deviated from the actual best dating hypothesis when they were not identical.

### Empirical data

We analyzed two empirical datasets to test the applicability of the λ(*t*)likelihood approach. The first dataset addressed relationships and estimated divergence times between beetle species of the supertribe Clavigeritae (Coleoptera: Staphylinidae: Pselaphinae) (Parker and Grimaldi, 2014). We calibrated the tree with a Bayesian approach using the software MCMCtree and all possible subsets of calibration points, implemented as soft minimum bounds. Subsequently, we compared the fit of these differently calibrated trees to the fossil record, using the empirical λ(*t*)derived from the PaleoDB data. The second empirical dataset we re-analyzed addressed insect inter-ordinal relationships (Misof *et al*., 2014). The original analysis of this dataset was based on separate tree calibrations for each meta-partition of the molecular data using identical tree topologies and fossil calibration points. These separate calibrations showed extensive variability of node age estimates. Here, we used our approach to check which of the separate calibrations best fit the fossil record. Since each calibration was based on an identical set of fossils and different sequence data meta-partitions, we were able to rank these calibrations by using the λ-score. We calculated the λ-score for a total of 105 meta-partitions using 28 fossil calibration points provided by Misof *et al*. (2014). We calculated λ-scores for three different calibration schemes proposed by Tong *et al*. (2015), which are based on the amino acid sequence data and meta-partition schemes published by Misof *et al*. (2014). Please note that Misof *et al*. (2014) applied log-normal priors on node age estimates, while Tong *et al*. (2015) applied in one of their analyses uniform priors with soft bounds and in another one soft minima. In all three calibration schemes, a total of 85 meta-partitions were considered. In our evaluation of the three calibration schemes, we excluded several calibration points used in the published study on insect relationships (Misof *et al*., 2014), because they were either beyond the current time range of the empirically inferred λ(*t*) (fossil IDs F1, F3, F35, and F36 listed in supplementary Table 8 published by Misof *et al*. (2014)) or node age estimates were beyond the range of λ(t) (fossil IDs F2, F4, F7, F10, F11, F16, F17, F31, and F33, supplementary Table 8 published by Misof *et al*. (2014)). Futhermore, we excluded the fossil calibration points with IDs F8, F9, and F24 (supplementary Table 8 published by Misof *et al*. (2014)), because a large number of node age estimates associated with this fossil were younger than the presumed fossil dating. A consensus dating hypothesis was inferred from the ten highest scoring meta-partitions by calculating the mean node ages.

## Acknowledgments

This work was supported by the Leibniz Association and was conducted within the Leibniz Graduate School on Genomic Biodiversity Research at the Zoological Research Museum A. Koenig and University of Bonn, Germany. We would like to express our gratitude to Jun Tong, Simon Ho, and Nathan Lo for making data available to us. We thank Duane McKenna for helpful comments on the manuscript. Furthermore, we thank all members of the Leibniz Graduate School for fruitful discussions.

